# Identification and Expression Analysis of CDPK Family in *Eriobotrya japonica*, reveals *EjCDPK25* in Response to Freezing Stress in Fruitlets

**DOI:** 10.1101/2024.05.01.591999

**Authors:** Yifan Xiong, Shunquan Lin, Jincheng Wu, Shoukai Lin

## Abstract

The fruitlets of loquat (*Eriobotrya japonica* Lindl.) are susceptible to freezing injury due to their developmental cycle encountering winter. Freezing stress severely damages the fruitlets, resulting in loss of fruit yield and quality. Studies have shown that Ca^2+^, as a second messenger, is involved in signal transduction in loquat fruitlets under freezing stress. However, the mechanism of downstream calcium signal transduction in loquat fruitlets under freezing stress is currently unclear. Calcium-depend protein kinase (CDPK) as the most particular calcium sensor family in plants, play an important role in multiple stress signal transduction including freezing. In this study, we identified the loquat CDPK family on a genome-wide scale. A total of 34 *EjCDPK* genes were identified and studied for basic structural and phylogenetic features. EjCDPKs can be divided into four subgroups phylogenetically. The patterns of exon-intron and protein motif are highly conserved among the subgroups. Collinearity analysis identified several segmental duplicate events in EjCDPK family. RNA-seq based transcription analysis indicated that partial of *EjCDPK*s differently expressed in response to freezing stress with tissue-specific. Moreover, we preformed correlation analysis between expression value and trait data of loquat fruitlet under freezing stress by weighted co-expression gene network. After that, *EjCDPK25* was selected as the candidate because of its potential freezing stress response function. Protein kinase related GO terms were enriched in *EjCDPK25* co-expression genes, and then QPCR was performed to examine the target gene’s expression pattern. In addition, *EjCDPK25* was cloned to construct overexpression vector to obtain transgenic *Arabidopsis* plants. Transgenic and wild-type *Arabidopsis* were suffered freezing stress treatments (-5°C). The results showed that the survival rate of *EjCDPK25* overexpressing transgenic *Arabidopsis* was significantly higher than WT. In summary, this study identified loquat CDPK family firstly, and our data provide significant insights into the evolution and function of loquat CDPKs. Above all, a freezing stress response gene *EjCDPK25* was verified can increase the resistance of freezing stress in *Arabidopsis*.

## 1. Introduction

Loquat (*Eriobotrya japonica* Lindl.) is a conventional commercial crop originated in China and then spread around the world, represents a sweet-acid fruit with special flavour^[1]^. Even as a subtropical evergreen fruit tree, the loquat has relatively strict requirements for the cultivation environment. The fruit development cycle of loquat always meets with winter and the fruitlets are susceptible to cold temperature. Frost can cause fatal damage to fruitlets and seriously threaten the production of loquat. Unless trunk can withstand temperatures from -12°C to -18.1°C and flower buds can tolerate temperatures below -6°C, loquat fruitlet are more sensitive to low temperature, easily damaged by frozen at -3°C^[2]^. In recent years, due to the earth’s climate anomalies, loquat freezing stress has occurred frequently, causing a large reduction in production and serious economic losses.

Under -3°C, the ultrastructure of loquat fruitlets shows that, protoplasmic membrane and vesicle membrane rupture, protoplasts concentrate, and chloroplasts were distorted and deformed, mitochondrial membrane structure was damaged, and the inner ridge was lost^[3]^. Loquat fruitlets infected with Ice Nucleation Bacteria (INA) were more sensitive to freezing stress, and can increased the damage more than 50%^[4]^. Trifluralazine, as the calmodulin-specific antagonist may regulate the AsA-GSH cycle in loquat fruitlet under low temperature stress by inhibiting the Ca^2+^-CaM signaling pathway^[5]^. Low temperature stress can causes a decrease in the amount of Ca^2+^ bound to the cell membrane of loquat fruitlet, induces an increase in the activity of lipid degrading enzymes and lipoxygenases, and reduces the structural stability of the cell membrane^[6]^. In loquat seedlings, exogenous Ca^2+^ increased the activity of Ca^2+^-ATPase on mitochondrial membrane, that maintained the Ca^2+^ signals in a low steady-state, and enhanced the activity of antioxidant system to reduce the low temperature damage^[7]^. Low temperature stress can also shut down the anti-oxidation system by reducing the activity of related enzymes such as glutathione peroxidase and glutathione-S-trasferase, while exacerbating the damage of membrane lipid peroxidation in loquat fruits^[8]^. Ca^2+^ can alleviates chilling injury in loquat fruit by regulating ROS homeostasis and maintaining membrane integrity ^[9, 10]^. Inspired by existing studies, Ca^2+^ came into our sight as an elixir to underlying the mechanisms of freezing stress signal transduction in loquat fruitlets. Plants have the ability to sense various stress signals from a changing environment and to transmit stress signals by multiple signal transduction mechanism in cells. Calcium (Ca^2+^) signaling is a prevalent pathway in plants with rapid response and high sensitivity^[11]^. Under normal conditions, the Ca^2+^ concentration in the cell maintaining a dynamic equilibrium, but under the stimulus of stress caused by the external environment, there is a rapid rise and fall in Ca^2+^ concentration, and finally a dynamic equilibrium is reached again^[12]^. Ca^2+^ channel proteins were anchored to cell membrane and pump free Ca^2+^ from extracellular into cell to generating cell-specific and stress-specific Ca^2+^ spikes through differentiated timing, intensity, and frequency^[13]^. These information can be decoded by calcium-binding protein, usually known as calcium sensors, to drive specific responses^[14]^. Large number and diversity of Ca^2+^-binding protein were found in plants, including a prototypical calcium sensor CAM/CML (Calmodulin and Calmodulin-like), Calcineurin B-like proteins (CBL) and Ca^2+^-dependent protein kinases (CDPK)^[15-17]^.

As a plant-specific multigene family, CDPKs exhibit distinct expression pattern and subcellular localization, playing versatile roles in activating and repressing of downstream substrate^[18]^. CDPKs have highly conserved protein structure, usually consist of four typical Ca^2+^-binding domain (EF-hand) at C-terminal and fused to a Ser/Thr kinase domain and a CDPK activation domain at variable N-terminal^[19]^. It is generally accepted that the activation of CDPK is controlled by pseudosubstrate mechanism, where structural changes allow the release of the pseudosubstrate from N-terminal kinase domain after EF-hands domain binding Ca^2+[20, 21]^. CDPKs are activated by Ca^2+^ binding and gain the ability to phosphorylate downstream targets and transduce Ca^2+^ signals into phosphorylation cascades^[22]^. All CDPKs have similar conserved molecular structures, however, some CDPKs show limited or no sensitivity to Ca^2+^ for their kinase activity ^[20]^. Therefore, the activation mechanism of CDPKs remains not fully understood.

CDPKs are widely identified in plants, there are 34 CDPKs in *Arabidopsis thaliana*^[23]^, 31 in rice (*Oryza sativa*)^[24]^, 35 in maize (*Zea mays*)^[25]^, 20 in wheat (*Triticum aestivum* L.)^[26]^, 19 in grape (*Vitis vinifera*)^[27]^ and 37 in apple (*Malus domestica*)^[28]^. Ample evidence shows that CDPK play crucial roles in plants abiotic stress response including cold, salt and drought stress^[29]^. In *Arabidopsis, AtCDPK10* was identified as an important regulatory component involved in drought stress response through stomatal movements modulated by ABA and Ca^2+^ signals^[30]^. Disrupted the expression of *AtCDPK23* can greatly enhanced *Arabidopsis* tolerance to salt and drought stress, however, over-expression *AtCDPK23* increased the plant sensitivity to salt and drought stress^[31]^.In rice, *OsCDPK7* was induced by cold and salt stresses, over-expression of *OsCDPK7* conferred both cold and salt/drought tolerance on rice plants and suppression of *OsCDPK7* expression lowered the stress tolerance^[32]^. *OsCPK17* is an indispensable response gene in cold stress, and likely affecting the activity of membrane channels and sugar metabolism^[33]^. OsCDPK24 phosphorylated downstream target OsGrx10 by controlling of calcium signal, and inhibit OsGrx10 activity to maintain high glutathione level to improve resistance of freezing stress in rice^[34]^. In maize, the expression of *ZmCPK1* can response to cold exposure, however, over-expression *ZmCPK1* reduce its resistance to cold stress indicate that *ZmCPK1* as a negative regulator of cold stress signaling^[35]^. Obviously, the identification and functional verification of CDPK family genes in crops and model plants have largely studied already.

According to the above, calcium signal that engaged in plant cold stress responses was widely proven. However, there are few studies to reveal the mechanisms of calcium single regulation in loquat fruitlet, especially absence of the identification of calcium sensor CDPK family and its mechanism study. In this study, loquat CDPK family was identified by genome-wide BLAST and domain motif scanning. After cold stress treatment of transgenic *EjCDPK25 Arabidopsis*, our result indicated that *EjCDPK25* is positively involved in cold stress response.

## 2. Materials and Methods

### 2.1 Identification of CDPK genes in *E*.*japonica*

To determine *CDPK* genes in *E*.*japonica*, the latest reference genome and annotation of ‘JieFangZhong’ loquat were obtained from CNGB(https://db.cngb.org/cnsa/), using the accession number of CNP0001531^[36]^. DNA and protein sequence of *Arabidopsis thaliana* and *Oryza sativa* CDPK family were download from Uniport (https://www.uniprot.org/). *Malus domestica* genome V3.0 was obtained from *Rosaceae* genome data base(www.rosaceae.org). According to the identified *MdCDPK* gene id^[28]^, extracted the sequence from *M*.*domestica* genome. *Vitis vinifera* genome and annotation were download from Grape Genome Database(http://www.genoscope.cns.fr/externe/GenomeBrowser/Vitis/). The sequences of *VvCDPK* were extracted by their gene id^[27]^. Totally, sequences of 34 *AtCDPK*, 31 *OsCDPK*, 37 *MdCDPK* and 19 *VvCDPK* were collected. Then, we downloaded the Hidden Markov Model (HMM) of EF-hand domain (PF13499) and protein kinase domain (PF00069) that are both indispensable to CDPK family. HMMER software^[37]^ was used to screen loquat protein sequences with EF-Hand domain and protein kinase domain with e-value set as 0.01. We also performed the local BLAST software^[38]^ to run BLASTP within CDPK sequence mentioned above with e-value less than e^-5^. Sequence similarity less than 50% were cutoff. After that, all candidates were verified by SMART and Pfam databases. Finally, the CDPK family in *E*.*japonica* were identified without redundant. Molecular weights (Mw) and isoelectric points (*pI*) of EjCDPKs were predicted by ExPASy (https://www.expasy.org/)^[39]^.

### 2.2 Chromosome localization and Phylogenetic analysis

Chromosome mapping of *EjCDPK* genes was accomplished by TBtools^[40]^ based on the start and end positions extracted from genome annotation. Multiple sequence alignment carried by ClustalW algorithm. And then, neighbor-joining (NJ) phylogenetic tree with 1000 bootstrap value was construct by MEGA v10. Software^[41]^. Moreover, an online software iTOL ([https://itol.embl.de/)^[42]^ was used to beautified the genetic tree.

### 2.3 Gene structure and protein motif analysis

By using the online software Gene Structure Display Server (GSDS) ([http://gsds.cbi.pku.edu.cn/)^[43]^, gene extron-intron patterns were determined. The MEME protein conserved domain analysis tool ([http://meme-suite.org/tools/meme)^[44]^ was used to analyze all the EjCDPK protein sequences with classic mode, and setting the maximum motif number set as 8.

### 2.4 Gene duplication and synteny analysis

MCScanX software^[45]^ was applied to identify the segmentally duplicate and tandemly duplicate of *EjCDPK* genes. Moreover, synteny analysis of *EjCDPK* genes between *A. thaliana* and *M. domestica* was also used MCScanX. And the result of duplication and synteny analysis was visualized by TBtools.

### 2.5 Analysis of cis-element in *EjCDPK* genes

5’ upstream 2000bp sequences of *EjCDPK* genes were extracted from loquat genome, and were submitted to PlantCARE ([https://bioinformatics.psb.ugent.be/webtools/plantcare/html/)^[46]^ for analysis. After analysis the cis-acting elements, abiotic stress response element was retained and visualized by TBtools.

### 2.6 Expression profile of *EjCDPK* genes in fruitlet under cold stress

RNA-seq raw data of 54 samples of ‘Zaozhong6’ loquat fruitlets under cold stress was obtained from our former research (Unpublished), including fruit and seed tissue of fruitlet treated at three times (2h, 4h and 6h) scales and three temperature (25°C, -1°C and -3°C) scales. Trimmomatic software (v.3.0)^[47]^ was used to filter out low-quality reads and trimming sequencing adaptors. Fastqc software was used to control the reads’ quality. After that, all clean reads were mapped to the *E*.*japonica* reference genome by Hisat2 (v.2.1.0)^[48]^. SAMtools software (v.1.4) ^[49]^ was used to convert the Sam format file into sorted Bam format. Cufflinks software (v.2.2.1)^[50]^ was applied to calculate the FPKM value of each sample and export the total expression matrix. TBtools was used to extract the expression matrix of *EjCDPK* genes and plot a heatmap with their FPKM values.

### 2.7 Weighted gene co-expression network construction and key *EjCDPK* gene select

Standardized expression data FPKM of 54 samples were used for WGCNA analysis. R software (v 4.11) and R package WGCNA (v 1.70.3)^[51]^ were applied for construct weighted gene co-expression network. MAD (median absolute deviation) was used to filter the input gene expression matrix, reserving only top 10,000 genes by sorting. Sample cluster should be carried out first, and the outlier samples should be removed to exclude their influence on the whole data. Data after removing outlier samples were used to calculate the soft threshold β for constructing the scale-free distribution network. After selecting suitable β value, TOM (Topological Overlap Matrix) is constructed from gene expression data using this threshold. Then, the TOM was clustered by hierarchical and constructed clustering tree. Branches of the hierarchical clustering tree are cut and distinguished, and the Cluster Dendrogram is finally obtained. Physiological and phenotypes of loquat fruitlet under cold stress, including fruit hardness, relative electrical conductivity (REC), malondialdehyde (MDA) and proline content, were correlated with weighted gene co-expression network.

### 2.8 GO and KEGG analysis of *EjCDPK25* co-expression genes

Annotations background for GO and KEGG of *E*.*japonica* were obtained from eggNOG annotate tool^[52]^ by uploading all *E*.*japonica* protein sequences. Annotations files were split by TBtools and use for enrichment backgrounds. Enrichment analysis was subjected to R package ClusterProfiler (v4.2.2)^[53]^.

### 2.9 Quantitative real-time PCR analysis of *EjCDPK25*

cDNA of 54 loquat fruitlet samples were used as the templet of qRT-PCR. The primers used for qRT-PCR were listed in supplement file. Bio-RAD CFX96 system was applied to perform qRT-PCR with TB Green Premix Ex Taq (Takara). Relative expression level calculation method was follow as described.

### 2.10 Vector construction and plant transformation

*EjCDPK25* cDNA was amplified by PCR and gel extraction. Then, *EjCDPK25* was cloned into *pCAMBIA1301* vector using In-Fusion HD Cloning Kit (Takara). After that, the constructed vector was transformed into *Agrobacterium* strain GV3101. In order to obtain the transgenic *Arabidopsis*, floral dip method was applied. Transgenic *Arabidopsis* were seeded on half-strength MS medium containing hygromycin B to perform select.

### 2.11 Plant material and growth condition

*Arabidopsis* seedings were grown in plant incubator at 22°C under 16h/8h light and dark conditions. The Petri dishes containing MS medium with 0.8% agar. Plants after seeding were growth in pot filled up with nutrient soil and vermiculite (3:1).

### 2.12 Cold tolerance treatment assays

2 weeks old transgenic *Arabidopsis* and wild type col-0 *Arabidopsis* were used for cold stress treatment. After set the plant incubator’s temperature as -5°C, use mercurial thermometer to supervise whether the temperature is stable. When the temperature is settled down, *Arabidopsis* on the Petri dishes were directly subjected to cold treatment, last for 3.5h. When finished cold stress treatment, the *Arabidopsis* were subjected to 4°C chamber recovery 12h under dark condition and then grown at normal condition for next 10 days. At last, survival rates were calculated by number of living plants divided number of total plants.

## 3. Results

### 3.1 Identification of CDPK genes in *E*.*japonica*

By combined utilization of HMMER and BLAST, totally 34 *EjCDPK* genes were identified in *E*.*japonica* genome-wide. Basic information including gene id, length of CDS and protein, pI and Mw were shown in Table 1. The protein length of EjCDPK ranges from 417 to 676 amino acids, with a theoretical isoelectric point of 5.12 to 9.23. Predicted molecular weight of EjCDPKs range from 47.89 to 76.07kDa, with an average 61.94 kDa.

### 3.2 Chromosome localization and phylogenetic analysis

*EjCDPK* genes distribute on 14 chromosomes of *E*.*japonica* genome, except for chromosome 1, 8, 13 and 16. Numbers of *EjCDPK* genes on each chromosome ranges from 1 to 4 (Figure 1). In addition, the names of 34 *EjCDPK* genes were determined by the localization on 13 chromosomes. Chromosome 11 and 14 only locate one *EjCDPK* gene, named *EjCDPK23* and *EjCDPK28* respectively. Chromosome 4, 6, 7 and 17 all locate two *EjCDPK* genes, listed as *EjCDPK8, EjCDPK9, EjCDPK13, EjCDPK14, EjCDPK15, EjCDPK16, EjCDPK33* and *EjCDPK34*. Chromosome 2, 5, 9 and 10 locate three *EjCDPK* genes, range from *EjCDPK1-3, EjCDPK10-12, EjCDPK17-19* and *EjCDPK20-22*. Chromosome 3, 12 and 15 each locate four EjCDPK genes, named as *EjCDPK4-7, EjCDPK24-27* and *EjCDPK29-32*. Neighbor-joining tree of CDPK genes in five species was constructed by MEGA and embellished by iTOL was shown in figure 2. *EjCDPK* genes can be divided into four subgroups, according to the distribution of CDPK family in *A. thaliana* and *O. sativa, M. domestica* and *V. vinifera*. Four subgroups named as EjCDPK I, EjCDPK II, EjCDPK III and EjCDPK IV, containing 13, 8, 10 and 3 *EjCDPK* genes respectively. *M. domestica* and *E*.*japonica* both belong to *Rosaceae*, however, the number of *CDPK* genes in these two species shows limited difference, and no obvious gene family expansion or contraction. Intriguingly, similar situation was detected in *A. thaliana* and *O. sativa*, except for *V. vinifera* who shows CDPK gene contraction among mentioned species.

**Figure 1.**
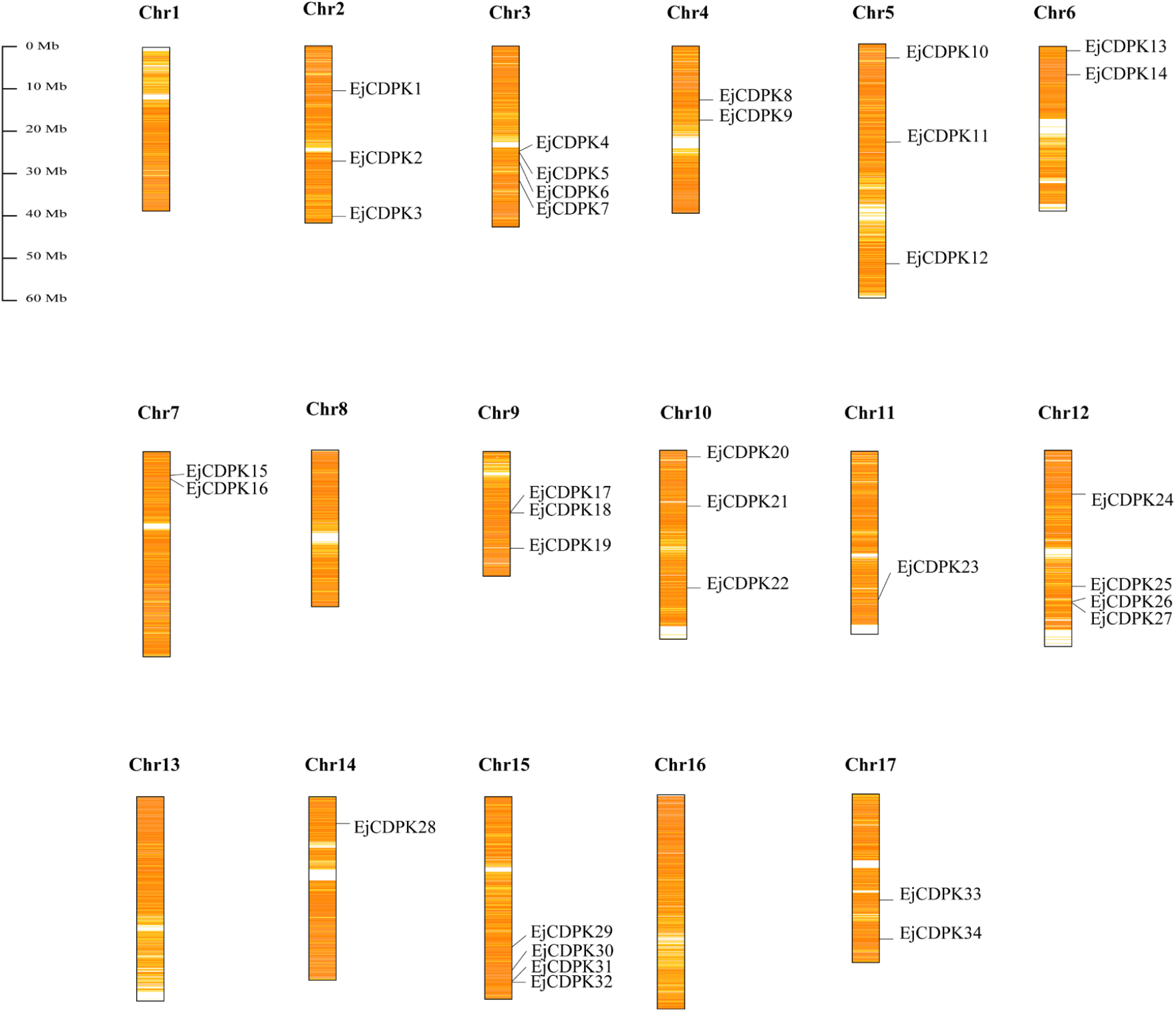
Chromosome localization of *EjCDPK*s. Rectangles represent loquat chromosomes that are drawn by scales. The internal filling heat map shows the gene distribution density on each chromosome. The specific location of EjCDPKs is indicated by the short black line.

**Figure 2.**
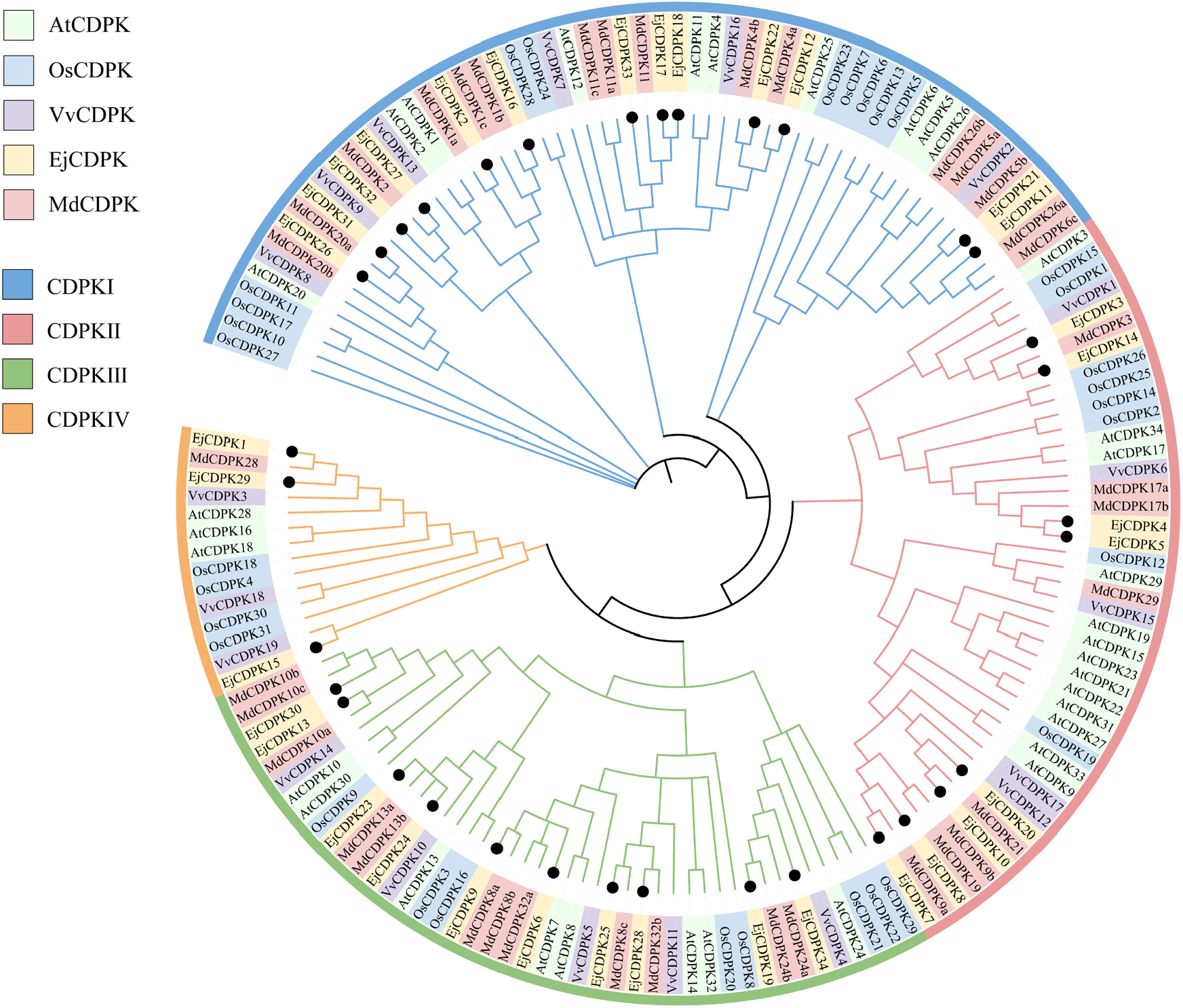
Phylogenetic analysis of CDPK family within loquat and other model plants. The full-length of amino acid sequence of CDPK from five species (*A. thaliana, O. sativa, M. domestica, V. vinifera* and *E*.*japonica*) were aligned by ClustalW. The phylogenetic tree was constructed using Neighbor-Joining method with 1000 bootstrap replicates by MEGA 10.0. CDPKs are shown in different colors represent five species, and the four subgroups are marked with distinct colors and Roman numerals I-IV.

### 3.3 Gene structure and protein motif analysis of EjCDPK family

In order to describe the structural diversity and evolutionary relationship between *EjCDPK* genes, we obtained coding sequence and full length of protein sequence of EjCDPK family. Intron-exon phase of *EjCDPK* genes were identified and visualized by GSDS tools. Protein conserved motifs were analyzed using MEME, the results were shown in figure 3. CDPK family members have more complicated function due to their distinguished structure among other two calcium sensor families. Protein kinase activity was enabled by Ef-hand domain capture calcium ions, and then CDPK catalyze downstream targets to transmit calcium ion signals. In EjCDPK I, *EjCDPK17, EjCDPK18* and *EjCDPK33* has no intron, and most other members has 7 exons. However, *EjCDPK2* and *EjCDPK26* has additional one and two exons, respectively. In EjCDPK II, *EjCDPK10* has 10 exons, *EjCDPK8* has 9 exons, and other 6 members all have 8 exons. Among EjCDPK III, most of the members have 8 exons, except *EjCDPK6* and *EjCDPK23* which has 9 exons and 7 exons. EjCDPK IV is the smallest subgroup, only have 3 members. *EjCDPK1* and *EjCDPK29* have the similar gene structure, both have 12 exons, while *EjCDPK15* has only 7 exons. The protein motifs of EjCDPK are highly similar, most EjCDPK have five protein kinase domains and two EF-Hand domains. However, members in EjCDPK IV and *EjCDPK24* which is belongs to EjCDPK III, are missing one EF-hand domain. And *EjCDPK8, EjCDPK34* and *EjCDPK15* all lacking one protein kinase domain. In general, EjCDPK family members shows highly similar and conservative in exon-intron phase and protein motif arrangement.

**Figure 3.**
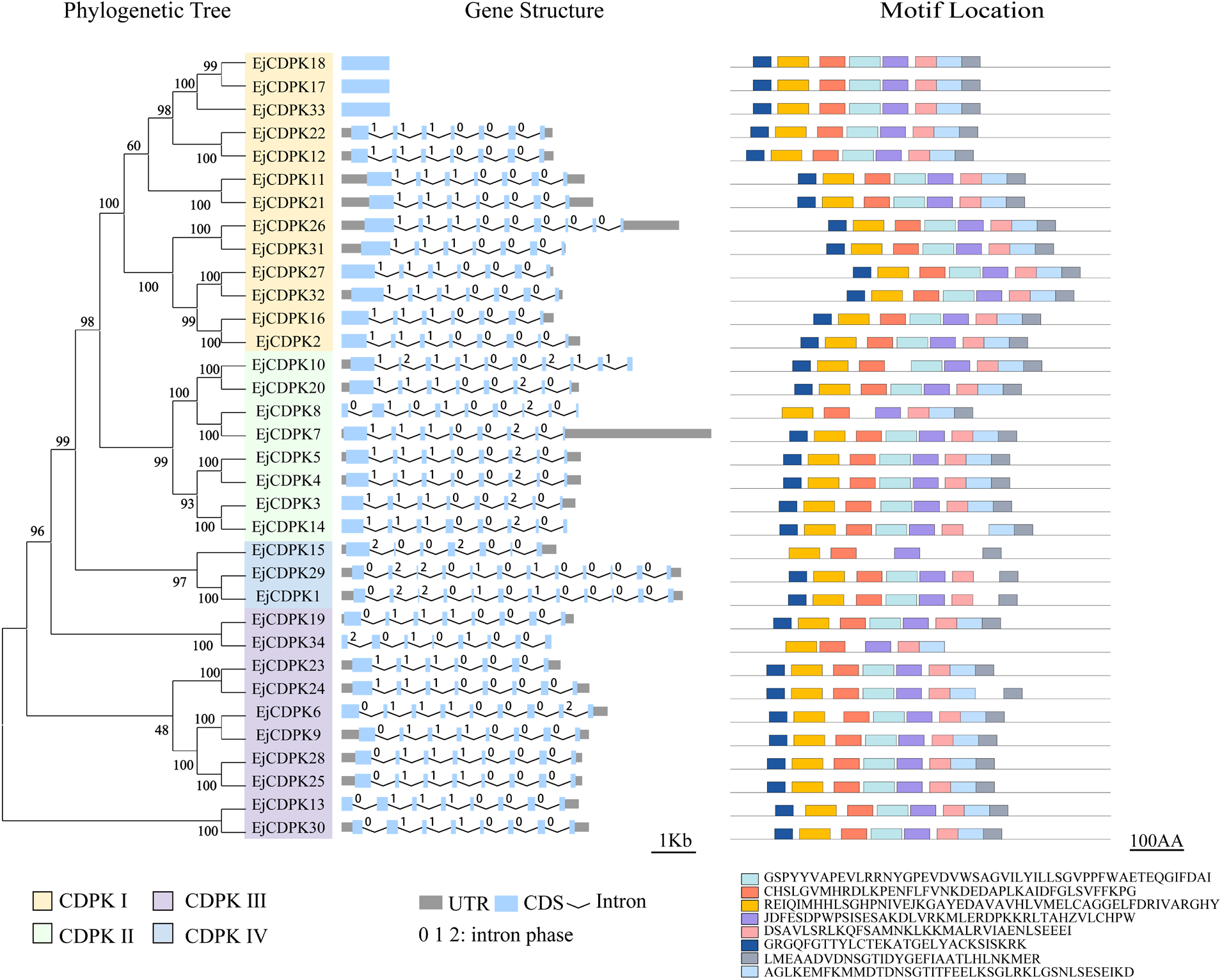
Gene structure and protein motif analysis of EjCDPKs. The unrooted phylogenetic tree was constructed by the use of full-length amino acid sequences of 34 *EjCDPK* genes with Neighbor-Joining method. Four subgroups are marked by distinct colors. (CDPK I yellow, CDPK II green, CDPK III purple, CDPK IV blue). The motif identification was used MEME online motif search tool by classic mode different motifs of respective EjCDPK are remarked by different colors and the consensus sequence of each motif was shown below the motif panel.

### 3.4 Duplication analysis of *EjCDPK* genes

Segmentally duplicate events and collinearity genes in EjCDPK family were identified by MCScanX, the result was shown in figure 4. Intriguingly, no tandemly duplicate of EjCDPK was detected in *E*.*japonica* genome. Totally 12 pair of collinearity genes were identified in *EjCDPK* genes. *EjCDPK7* and *EjCDPK8* has the same two collinearity genes *EjCDPK10* and *EjCDPK20*. Remaining 8 pairs of collinear genes were respectively are *EjCDPK2*/*EjCDPK16, EjCDPK3*/*EjCDPK14, EjCDPK6*/*EjCDPK9, EjCDPK11*/*EjCDPK21, EjCDPK17*/*EjCDPK33, EjCDPK25*/*EjCDPK28, EjCDPK26*/*EjCDPK31* and *EjCDPK27*/*EjCDPK32*.

**Figure 4.**
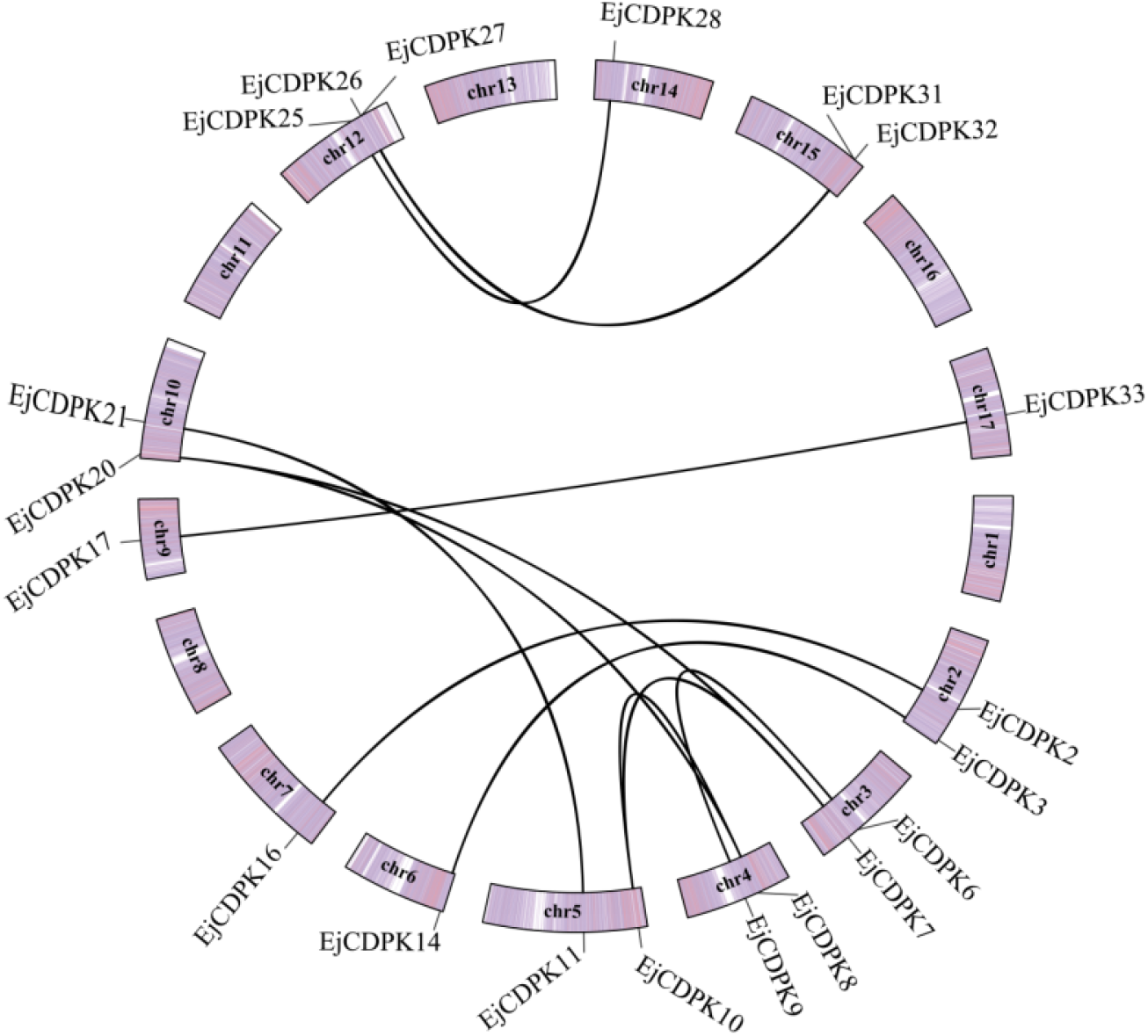
Synteny analysis in EjCDPK genes. The schematic diagram of 17 chromosomes of loquat is arranged in the form of circle, with the gene distribution and density heat map filled inside. The line between two genes on a chromosome indicates that this pair of genes have collinearity, and the short black line indicates the location of the gene on the chromosome.

Furthermore, collinearity analysis was also applied in *E*.*japonica* and other two species *A. thaliana* and *M. domestica*, results were shown in figure 5. As a result, 38 *EjCDPK* collinearity genes were identified in *A. thaliana* genome that distributed in five chromosomes. However, we detected 69 collinearity genes of *EjCDPK* in *M. domestica*, the number shows largely different compare to *A. thaliana*. Moreover, *M. domestica* chromosome 8, 14 and 16 has no *EjCDPK* collinearity genes.

**Figure 5.**
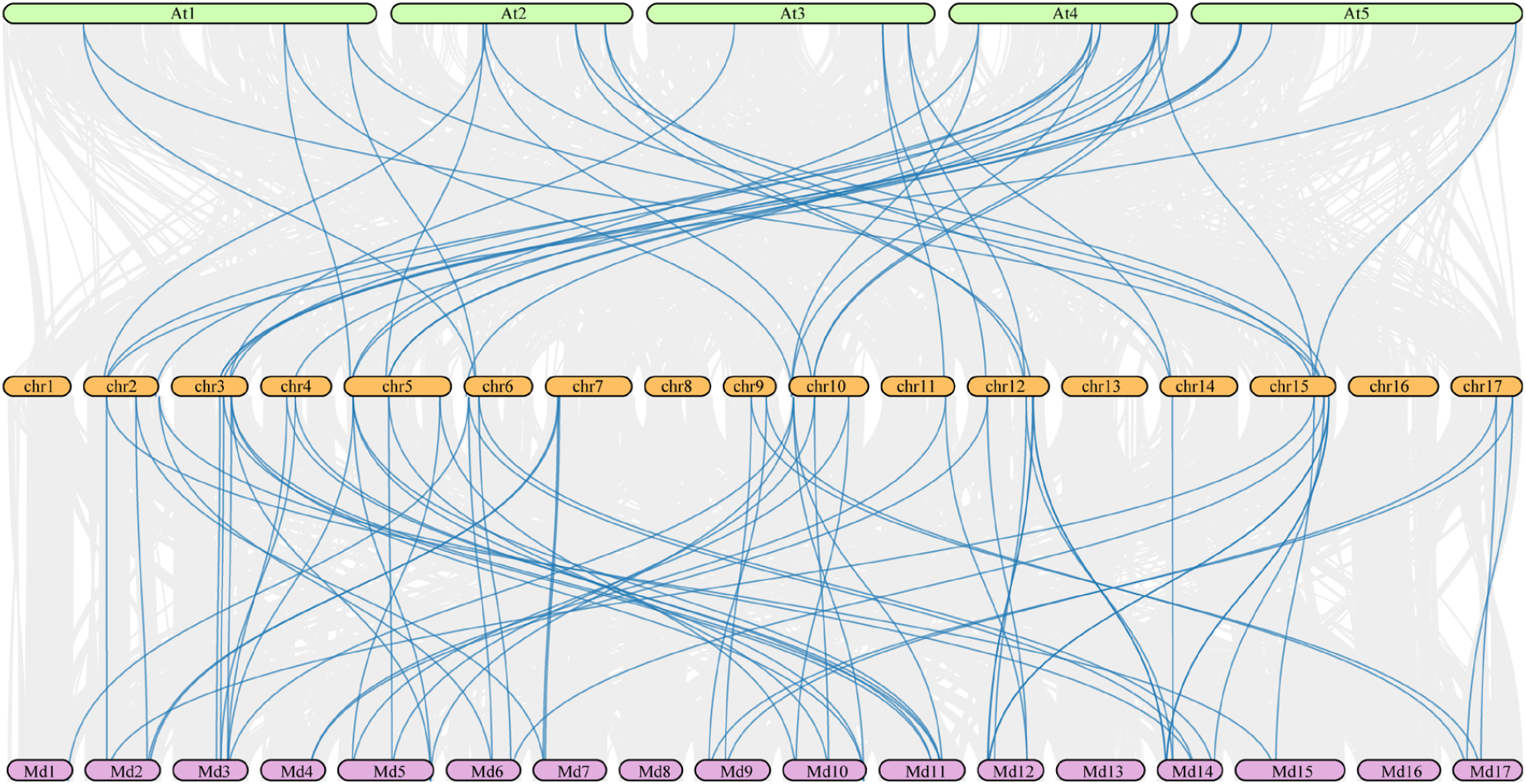
Synteny analysis among loquat, Arabidopsis and apple CDPK genes. Rectangle form with serial number represent the chromosomes of these three species, and were depicted in green, orange and pink. The approximate distribution of each *AtCDPK, EjCDPK* and *MdCDPK* is marked on the rectangle. Blue curves denote the syntenic gene pair.

**Figure 6.**
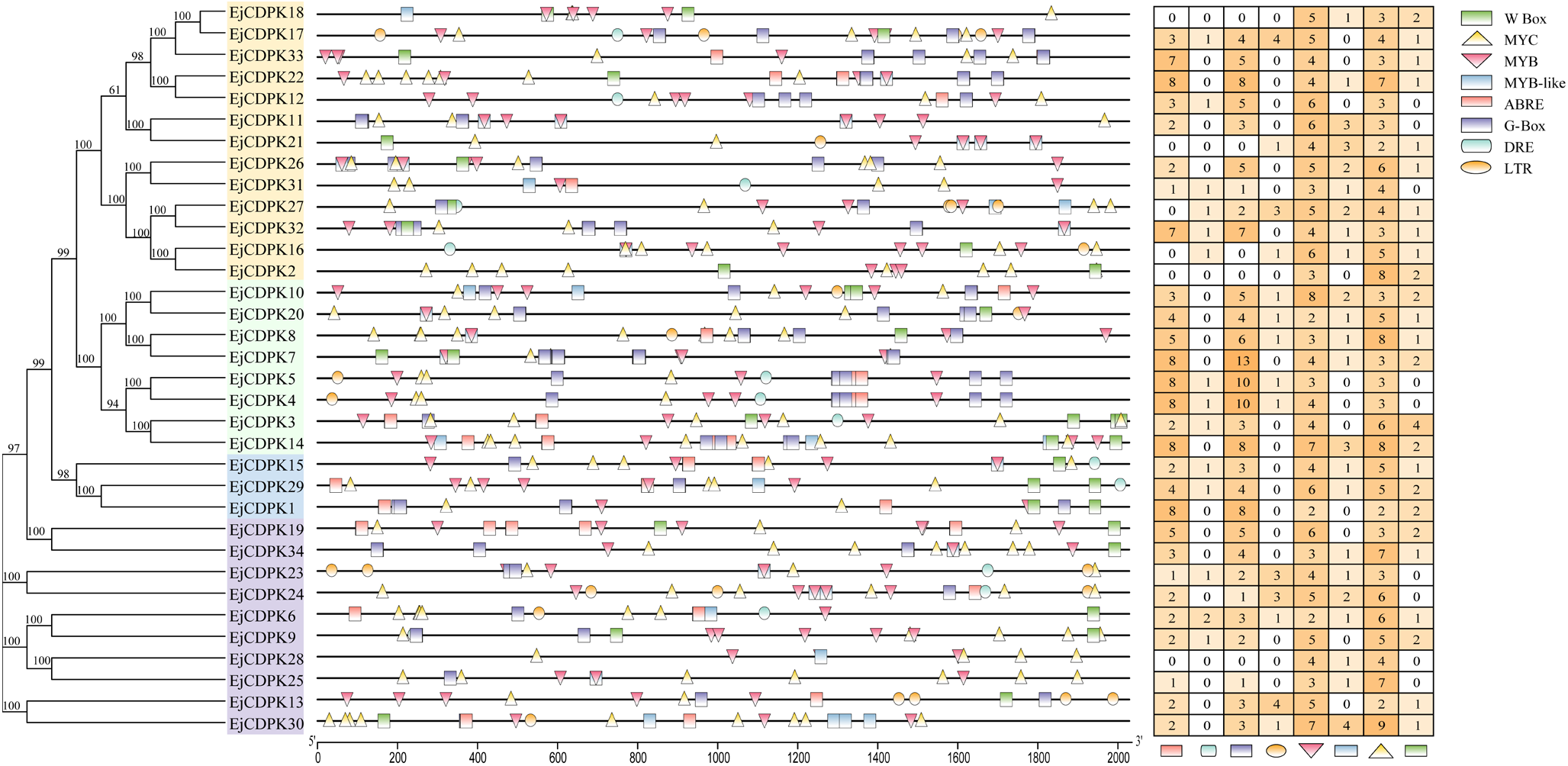
Stress related cis-acting element in the promoter region of EjCDPKs. Unrooted phylogenetic tree was constructed by the use of full-length amino acid sequences of 34 *EjCDPK* genes with Neighbor-Joining method. The location of each cis-acting element was shown on the line which indicate the 5’ upstream sequence of *EjCDPK*s by different shape. And the cis-acting element number of each EjCDPK was shown as heatmap.

### 3.5 Cis-element analysis of *EjCDPK* genes

Upstream 2000 bp region of *EjCDPK* genes were extracted to subjected to cis-element identification and analysis by PlantCARE. Abiotic stress response elements were selected especially cold response elements including DRE element, MYB element, MYB-like element, MYC element and LTR element. In EjCDPK family, 17.6% of the members lacking ABRE element (ABA-response element), EjCDPK II subgroup members has more ABRE elements than other subgroups. 58.8% of *EjCDPK* genes lacking DRE element, *EjCDPK6* has 2 DRE element which is the most. G-box element shows largely remained in *EjCDPK* genes, 85.3% of the member containing this element in their promoter region. Particularly *EjCDPK7*, which has 13 G-box elements. 41.2% of *EjCDPK* genes has LTR (Low temperature response) element, *EjCDPK13* and *EjCDPK17* both has 4. All the *EjCDPK* genes has MYB element and MYC element, however, only 67.6% has MYB-like element. 73.5% of *EjCDPK* genes has W-box element, and *EjCDPK*3 has 4 which is the most.

### 3.6 Expression profiles of *EjCDPK* genes under cold stress

Normalized gene expression value FPKM was counted by Cufflinks software (Figure 7). Expression fold change >2.0 was considered as differential expression. After clustering the expression profiles of *EjCDPK* genes by samples and treatment temperature, we found that differential expression of *EjCDPK* genes caused by treatment temperature is more obvious than tissue specific expression. 38.2% of EjCDPK genes shows lower expression, and no differential expression. Some of the *EjCDPK* genes have differential expression in different tissue. *EjCDPK25* and *EjCDPK28* were both up regulated response to -3°C treatment for 2h, 4h and 6h in loquat seed, and have differential expression. *EjCDPK16* was up regulated in loquat fruit under -1°C and -3°C treatment, however, only shows differential expression response to -3°C treatment. *EjCDPK7* and *EjCDPK17* were up regulated by -3°C treatment for 6h in loquat fruit, and shows differential expression. Furthermore, *EjCDPK29* were up regulated in both fruit and seed, shows differential expression. In loquat fruit, *EjCDPK29* was differential expressed after -3°C treatment for 6h. In loquat seed, unlike in fruit, *EjCDPK29* was up regulated in gradients of time, including 2h, 4h and 6h, and all shows differential expression.

**Figure 7.**
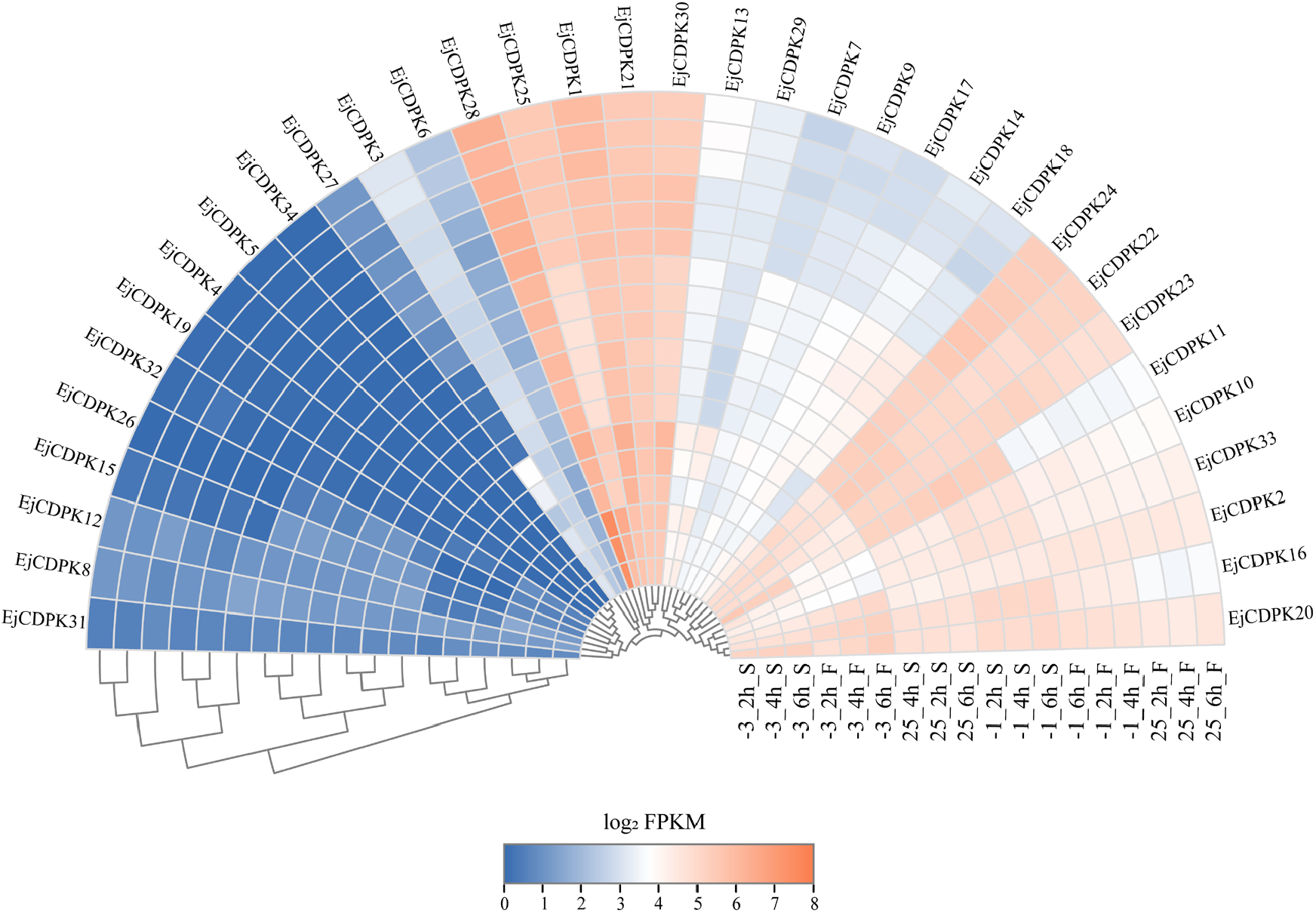
Expression patterns of *EjCDPK*s in loquat fruitlet under freezing stress. Base 2 logarithm of FPKM value was used to construct the heatmap. Freezing stress treatments including three temperatures (25°C, -1°C and -3°C), three gradients of time (2 hours, 4 hours and 6hours) and two tissues (S, seed and F, fruit).

### 3.7 Weighted gene co-expression network construction and key *EjCDPK* gene select

In order to narrow the range of target gene for functional verification, weighted gene co-expression network was constructed by R package WGCNA. Then the co-expression network was associated with loquat fruitlet trait data including hardness, relative electrical conductivity (REC), malondialdehyde (MDA) and proline content. SampleTree function was applied to find outlier samples, and no outlier was found in loquat fruit samples. Only one outlier, S163 was found in loquat seed sample. After cut off outlier, the expression matrix was subjected to calculate the soft threshold β. In loquat fruit samples, β was selected as 18, and selected as 7 in seed samples. Then weighted co-expression gene network in loquat fruit and seed were constructed by input β values respectively (Figure 8). 6 co-expression gene modules were clustered in loquat fruit expression data and 15 were clustered in seed expression data. After the construction of the weighted co-expression gene network, correlation analysis was conducted between the loquat fruitlet trait data and the co-expression network.

**Figure 8.**
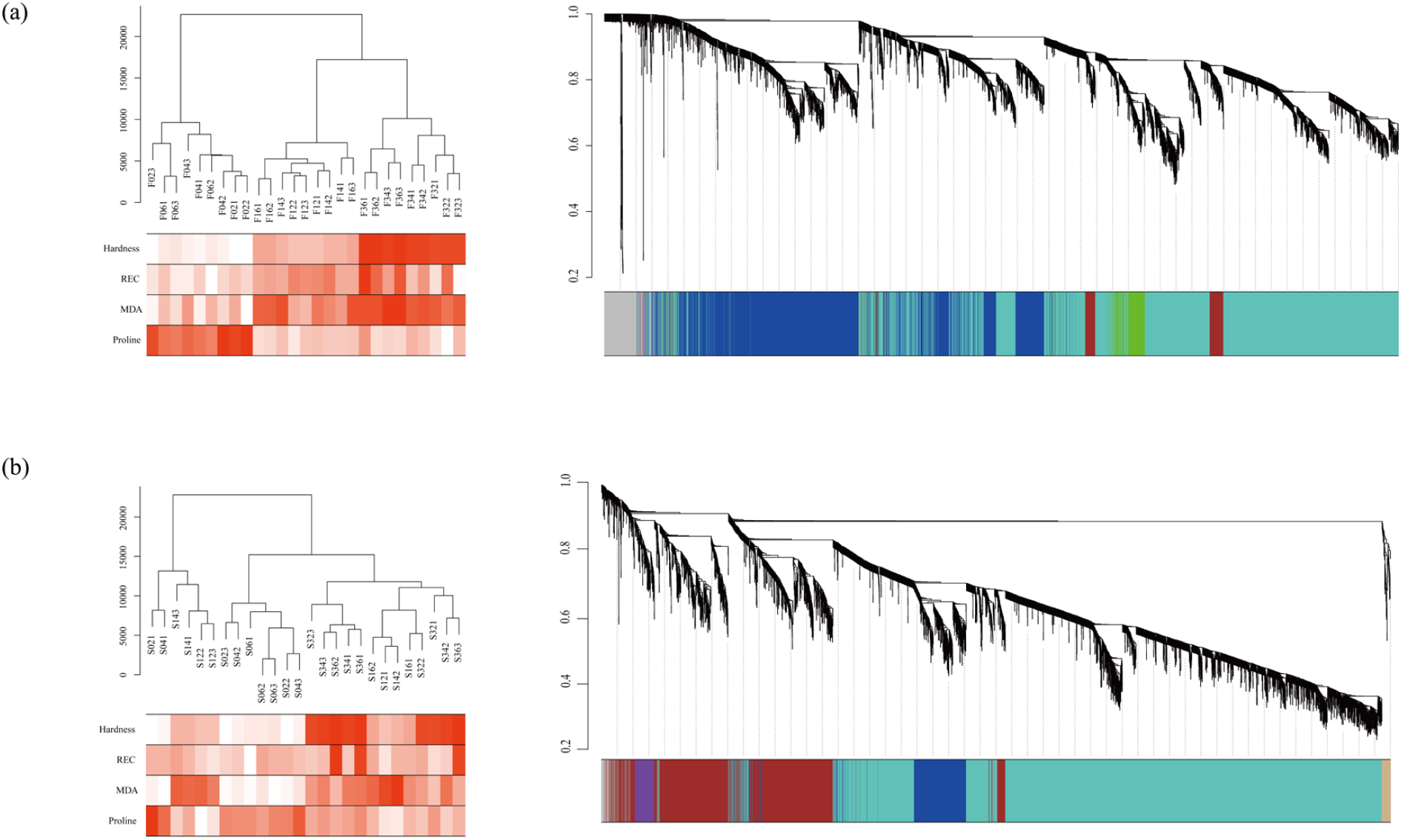
WGCNA by RNA-seq data form loquat fruit and seed under freezing stress. The left part of the figure shows the RNA-seq data sample cluster after cutoff outliers. And the relationship between sample expression and trait data. The soft threshold β for constructing TOM (Topological Overlap Matrix) was selected by set the independence corresponds as 0.8. The hierarchical clustering and module differentiation among genes are shown on the right. Genes with similar expression patterns belong to a branch, and different branches are cut and divided into different modules, which are represented by different colors.

**Figure 9.**
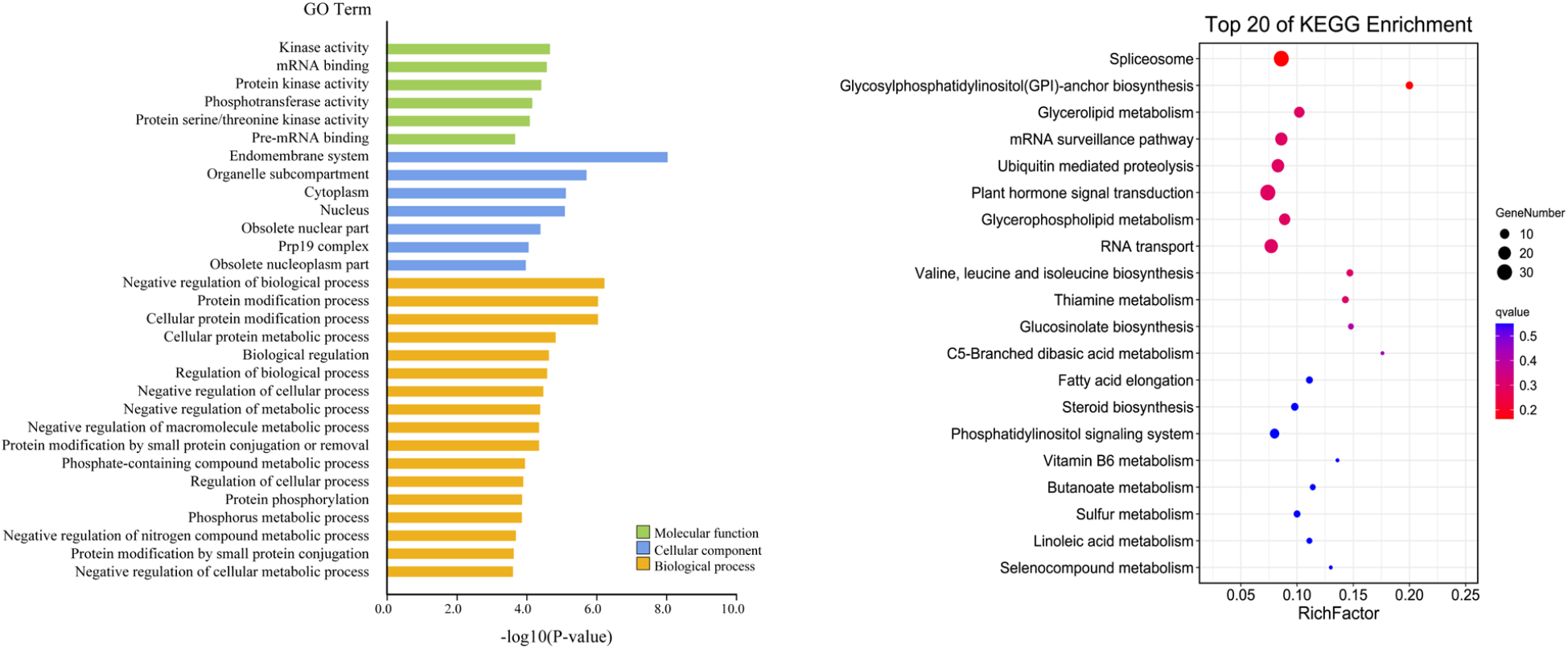
GO and KEGG enrichment of EjCDPK25 co-expression genes. Histogram shows the results of GO enrichment, three catalog of GO annotation was distinguished by different colors. Bubble diagram shows the results of KEGG enrichment.

Gene modules that have highly correlation (correlation coefficients >0.9) with loquat trait data were selected. As a result, turquoise module was selected not only in loquat fruit expression data (correlation coefficients is 0.96) but also in seed expression data (correlation coefficients is 0.93). Intriguingly, these two turquoise gene modules are both correlated with hardness. Moreover, eigengene expression patterns of turquoise modules were shown in figure 8. Then, we selected member relationship and gene significance both > 0.8 as threshold to filtered out the key genes in turquoise module, and picked up *EjCDPK* genes from them. As a result, *EjCDPK25* was came insight from loquat seed turquoise gene module. qRT-PCR was applied to verify the relative expression of RNA-seq data, and the result was shown in figure 10. The relative expression fold of *EjCDPK25* in loquat fruit was lower in the 2 hours at -1°C treated samples than the RNA-seq data, and the trend was similar in other treated samples. In loquat seed, the trend of relative expression fold of *EjCDPK25* was similar to RNA-seq data. And the relative expression fold of *EjCDPK25* in -3°C treated for 2, 4 and 6 h samples shown significant difference (P <0.05).

**Figure 10.**
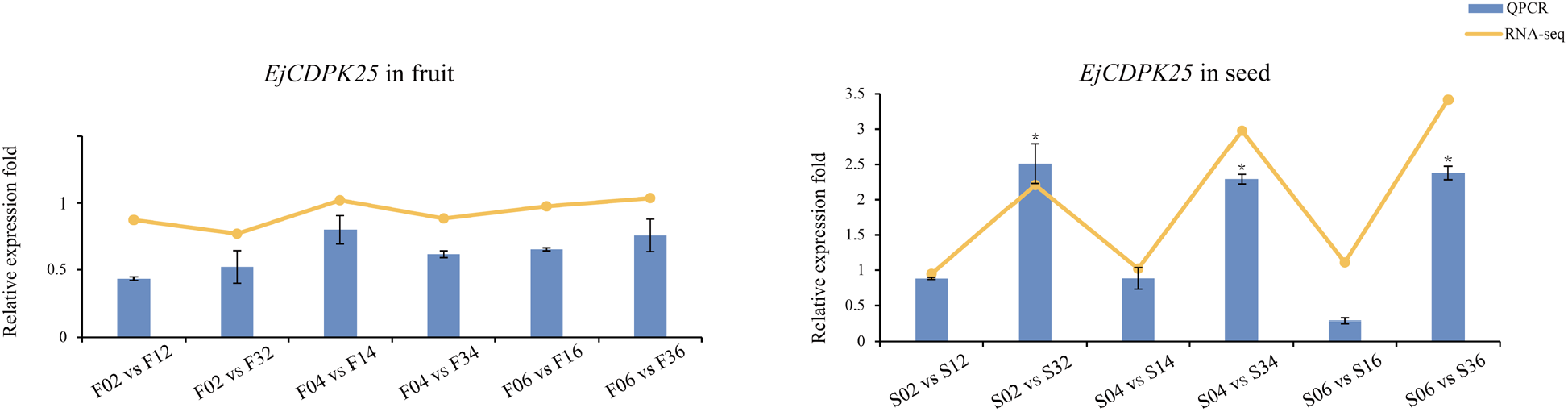
QPCR verification of EjCDPK25 gene’s expression. Y-axis represents the relative expression level of genes, and the X-axis represents the different treatment. Two tissues shown as F and S. 0, 1 and 3 were represent 25°C, -1°C and -3°C. 2, 4 and 6 were time gradients. The histogram with error bar represents the QPCR data, and the error bars were adding by standard error values (SEM). Line graph with endpoints represents RNA-seq data. Asterisk mark represents that the expression level of genes in the treated group was significantly higher than that in the control group (*:P<0.05).

**Figure 12.**
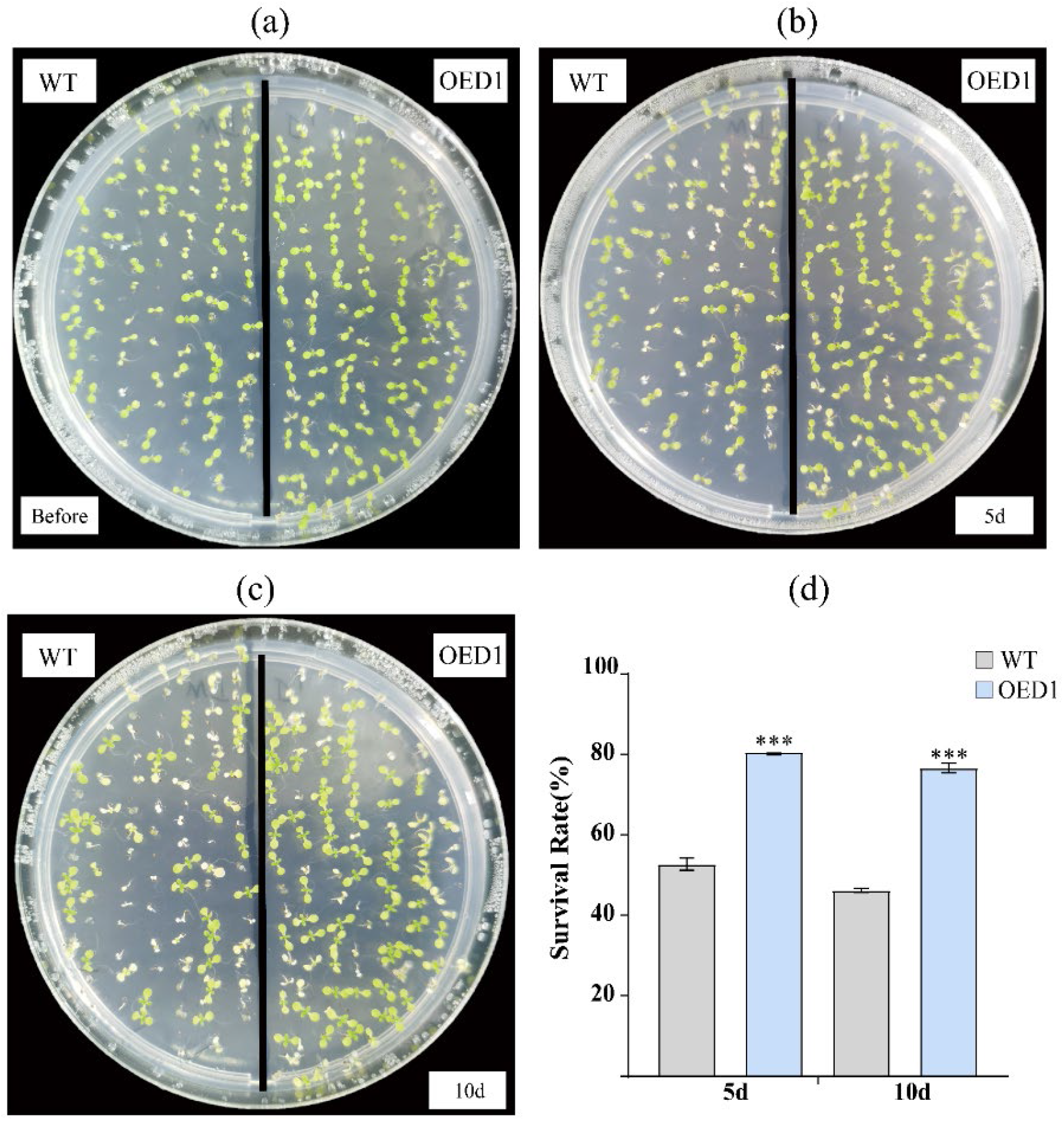
Transgenic Arabidopsis trait and survival rate under frezzing stress. (a), (b) and (c) shown the phenotypes of *Arabidopsis* under freezing stress treatments.5d and 10d represent recover from freezing stress for 5 days and 10 days. WT represents wild-type *Arabidopsis* and OED1 represents overexpressed *EjCDPK25 Arabidopsis*. (d) shows the survival rate of *Arabidopsis* after freezing stress treatment, and the error bar was added by standard error value (SEM). Asterisk indicates that the survival rate of transgenic *Arabidopsis* is significantly higher than the wild-type (***: P<0.01).

### 3.8 GO and KEGG analysis of *EjCDPK25* co-expression genes

The results of GO and KEGG enrichment analysis were shown in figure 9. In GO enrichment, a large number of protein kinase related items were concentrated in molecular functions category, including GO:0016301(kinase activity), GO:0004672(protein kinase activity), GO:0016773(phosphotransferase activity) and GO:0004674(protein serine/threonine kinase activity). Molecular functions category also contains mRNA binding related functional items such as GO:0003729(mRNA binding) and GO:0036002(mRNA precursor binding). GO:0012505(inner cell system), GO:0005737(cytoplasm) and GO:0005634(nucleus) were enriched in the cell component category. The co-expressed genes of *EjCDPK25* involved in biological processes include GO:0048519(negative regulation of biological processes), GO:0036211(protein modification), GO:0006468(protein phosphorylation) and GO:0031098(stress related protein kinase signaling cascade). In KEGG enrichment analysis, *EjCDPK25* co-expression genes were found to be enriched in Spliceosome, Glycerolipid metabolism, Ubiquitin mediated proteolysis and Plant hormone Signal transduction), etc.

### 3.9 Vector construction and *A. thaliana* transformation

The expression of *EjCDPK25* gene was verified by QPCR (Figure 10) and amplified by gradient PCR from loquat cDNA, shown in supplementary (Figure 2).

Then, amplified PCR products were cloned to T/A vector (pMD-18T, Takara). After sequenced, the target gene with In-fusion designed adapter were PCR-amplified using primer with restriction enzyme sites. PCR-amplified products were cloned into pCAMBIA1301 vector using In-fusion HD cloning kit (Takara). The confirmed clones by sequencing were applied to transformed *Agrobacterium* strain GV3101. (supplement file)

### 3.10 Cold stress treatment assays of overexpression *EjCDPK25 A. thaliana*

The 10-days-old *Arabidopsis* were treated at -5°C, and the survival rate of *Arabidopsis* was counted at 5 days and 10 days after treated recovery, results were shown in Figure 14. Three replicates were set up for cold stress. It was observed that part of the *Arabidopsis* was affected by cold stress, resulting in albinism and browning of leaves, and after few days the plant was died. For recovery 5 days, the mean survival rate of wild-type *Arabidopsis* was 52.8%, however, 80.5% for *EjCDPK25* overexpressed *Arabidopsis*, and shown significant difference (P <0.01). With increasing the recovery time under normal conditions, the phenotype of *Arabidopsis* damaged by cold stress became more obvious. After 10 days recovery, the mean survival rate of wild-type *Arabidopsis* decreased to 46.3%, and that transgenic *Arabidopsis* decreased to 76.6%. These results suggest that overexpression of *EjCDPK25* in *Arabidopsis* can promote the resistance to cold stress.

## 4. Discussion

Freezing stress threatened to the loquat fruits production severely. Especially in southeast China, where usually cultivated loquat varieties with excellent fruit quality but lower freezing stress resistance. ‘Zaozhong6’ loquat is one of the typical varieties. However, it was remained a huge obstacle to reveal freezing stress response mechanisms because of lacking the high-quality reference genome of loquat. In this study, we used the newest loquat reference genome, and applied both sequence homology and functional domain conservative methods to identify CDPK family. Totally 34 putative *EjCDPK* genes were identified and verified, excluded any redundant. EjCDPK and be divided into four subgroups according to the protein sequence similarity of AtCDPK (Figure 2). Intron-exon phase of *EjCDPK* genes is well conserved (Figure 3). The majority of *EjCDPK* genes containing seven or eight exons. The protein motifs of EjCDPK are also highly conserved. Most of EjCDPK has 2 EF-hands and 5 protein kinase domains. These data all indicate that, *EjCDPK* genes were derived from common ancestor via gene duplication as described in other species including *Arabidopsis* and rice^[54, 55]^. Segmental duplication and tandem duplication are two general formations of gene family^[56]^. Intriguingly, 12 *EjCDPK* collinearity genes were generated by segmental duplication events in loquat genome. But no tandem duplication was detected in *EjCDPK* (Figure 4). And the collinearity genes between loquat and apple are more than in *Arabidopsis* (Figure 5). Several cold stress response cis-elements were found in EjCDPK promoter region, like DRE, MYB, MYB-like, MYC and LTR (Figure 6). Among these elements, MYB, MYB-like and W-box are highly concerned due to their related transcription factors, which were found differential expressed under freezing stress in loquat fruitlets^[57]^. In *Arabidopsis*, the MYB15 protein was found interact with ICE1 and binds to MYB element in the promoters of CBF gene, negative regulate its expression in cold stress^[58]^. WRKY transcription factors are widely involved in abiotic stress responses of plants, and the W-box element is the binding site of WRKY. In the study of *AtPNP* gene promoter, W-box element was found indirectly regulated by salicylic acid and enhances abiotic stress resistance of *Arabidopsis*^[59]^. After mutated the core sequence of LTR element in barley, the resistance to low temperature stress was decreased, indicated that LTR element was involved in low temperature stress^[60]^. MYB, MYB-like and MYC element were all detected in *EjCDPK25* promoter region.

Previous observation found that with the increasing of freezing time at -1°C treatment, the browning of loquat fruitlet seed became more serious, but the change of fruit was not obvious. Neither fruit or seed can survive from -3°C treatment for 4 hours. Treatment at -3°C for 6 hours, loquat fruitlet was severely damaged and totally turned brown. After detected the expression pattern of *EjCDPK* genes, WGCNA correlated with loquat fruitlet trait data was performed to narrow the scale of candidates *EjCDPK* genes. Trait data including fruit hardness, relative electrical conductivity (REC), malondialdehyde (MDA) and proline content. During low temperature storage of loquat fruit, the increasing content of lignin and cellulose leads to the continuous increase of fruit hardness, severe damaged the fruit quality^[61]^. It has been reported that REC, MDA and proline are the indices of cold resistance^[62]^. By correlation analysis of WGCNA and trait data, turquoise module was picked up by its high correlation coefficients with fruit hardness both in loquat fruit and seed. After inner module selected, *EjCDPK25* gene came into our sight by setting the threshold above 0.9 both in gene significance and module membership (Figure 8). QPCR data of *EjCDPK25* in loquat seed shows the consistent expression trend with RNA-seq (Figure 10). Coexpression genes of *EjCDPK25* in loquat seed turquoise module were required for the GO and KEGG annotation to detected their functions (Figure 8). The majority annotated term of co-expression genes of *EjCDPK25* shows as protein kinase related, including GO:0016301 (kinase activity), GO:0004672 (protein kinase activity), GO:0016773 (phosphotransferase activity). Therefore, it is speculated that the co-expressed genes of *EjCDPK25* including the downstream targets to transmit the signals of low temperature stress. Moreover, it is also indicated *EjCDPK25* as calcium sensor maybe act as signal transmission hub under freezing stress^[63]^.

*EjCDPK25* was cloned and subjected to construct overexpression vector. *Arabidopsis* transformation by floral dip method, and T2 generation transgenic *Arabidopsis* were obtained. The survival rate of transgenic *Arabidopsis* significantly increased than wild type (Figure 14). This result speculated that *EjCDPK25* gene can enhanced the resistance to freezing stress in *Arabidopsis*. In *EjCDPK25* downstream regulation region, MYB and MYC cis-element were found. Further research is required to determine which transcription factors can regulate its expression under freezing stress.

Existing studies shown that, both positive regulation and negative regulation were found in plant CDPKs response to abiotic stress^[34, 35]^. However, in this study, we only focused on positive regulation of EjCDPKs in freezing stress. The down-regulated expression *EjCDPK*s and negative correlated gene modules in WGCNA were no further research. Subsequent studies could focus on this aspect of inference. The further studies of EjCDPK25 need to determine the downstream targets of EjCDPK25 by protein-protein interaction analysis. And homology overexpression of EjCDPK25 can provide stronger evidence than transgenic *Arabidopsis*. These intensive studies can draw a bigger picture of *EjCDPK25* regulation network under freezing stress in loquat.

In summary, our study firstly identified the CDPK family in loquat, and confirming that *EjCDPK25* could enhance the freezing stress resistance in *Arabidopsis*. This study can provide new insights for the freezing stress response mechanism of young loquat fruit.

