## Supplemental table and figures for "Identification and Expression Analysis of CDPK Family in *Eriobotrya japonica*, reveals *EjCDPK25* in Response to Freezing Stress in Fruitlets"

### SUPPLEMENTARY MATERIALS

### Supplementary table and figures

#### Table S1 Basic characteristic and physicochemical properties of *E.japonica* CDPKs

| Gene name | Chr | Group | CDS (bp) | Protein Length (aa) | pI | MW (kDa) |
|---|---|---|---|---|---|---|
| *EjCDPK1* | 2 | IV | 1668 | 555 | 9.23 | 63.17 |
| *EjCDPK2* | 2 | I | 1791 | 596 | 5.39 | 66.17 |
| *EjCDPK3* | 2 | II | 1602 | 533 | 6.09 | 59.76 |
| *EjCDPK4* | 3 | II | 1605 | 534 | 5.67 | 59.76 |
| *EjCDPK5* | 3 | II | 1605 | 534 | 5.67 | 59.76 |
| *EjCDPK6* | 3 | III | 1803 | 600 | 6.28 | 66.97 |
| *EjCDPK7* | 3 | II | 1617 | 538 | 7.26 | 60.61 |
| *EjCDPK8* | 4 | II | 1341 | 446 | 6.69 | 50.55 |
| *EjCDPK9* | 4 | III | 1605 | 534 | 6.11 | 60.07 |
| *EjCDPK10* | 5 | II | 1992 | 663 | 6.57 | 73.87 |
| *EjCDPK11* | 5 | I | 1716 | 571 | 6.04 | 64.00 |
| *EjCDPK12* | 5 | I | 1488 | 495 | 5.36 | 55.66 |
| *EjCDPK13* | 6 | III | 1677 | 558 | 6.6 | 62.98 |
| *EjCDPK14* | 6 | II | 1746 | 581 | 6.02 | 65.26 |
| *EjCDPK15* | 7 | IV | 1581 | 526 | 5.67 | 58.70 |
| *EjCDPK16* | 7 | I | 1764 | 587 | 5.43 | 65.22 |
| *EjCDPK17* | 9 | I | 1497 | 498 | 5.12 | 55.99 |
| *EjCDPK18* | 9 | I | 1497 | 498 | 5.16 | 55.59 |
| *EjCDPK19* | 9 | III | 1614 | 537 | 5.32 | 61.21 |
| *EjCDPK20* | 10 | II | 1641 | 546 | 6.45 | 61.11 |
| *EjCDPK21* | 10 | I | 1713 | 570 | 5.85 | 63.86 |
| *EjCDPK22* | 10 | I | 1503 | 500 | 5.42 | 56.17 |
| *EjCDPK23* | 11 | III | 1584 | 527 | 5.95 | 59.41 |
| *EjCDPK24* | 12 | III | 1722 | 573 | 6.16 | 64.20 |
| *EjCDPK25* | 12 | III | 1596 | 531 | 5.92 | 59.82 |
| *EjCDPK26* | 12 | I | 2031 | 676 | 5.85 | 76.07 |
| *EjCDPK27* | 12 | I | 1974 | 657 | 5.98 | 73.44 |
| *EjCDPK28* | 14 | III | 1596 | 531 | 6.16 | 59.98 |
| *EjCDPK29* | 15 | IV | 1629 | 542 | 8.9 | 61.12 |
| *EjCDPK30* | 15 | III | 1647 | 548 | 6.7 | 62.22 |
| *EjCDPK31* | 15 | I | 1815 | 604 | 5.85 | 67.21 |
| *EjCDPK32* | 15 | I | 1944 | 647 | 5.91 | 71.93 |
| *EjCDPK33* | 17 | I | 1509 | 502 | 5.22 | 56.13 |
| *EjCDPK34* | 17 | III | 1254 | 417 | 5.61 | 47.89 |

(a) (b)

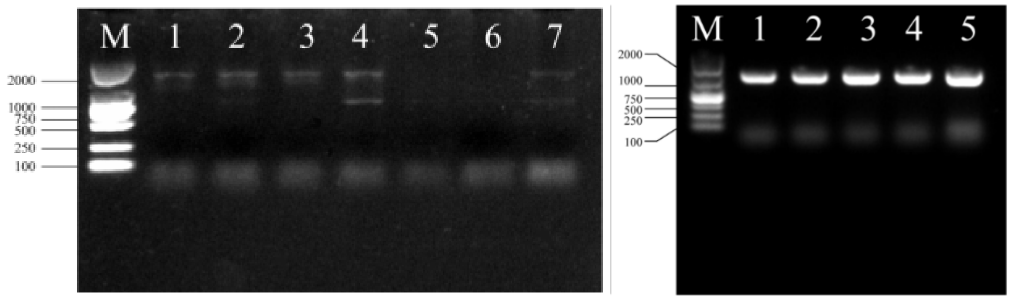


**Figure S1. Over-expression vector construction of EjCDPK25**

(a) Cloning of *EjCDPK25*. M represents DL2000 DNA marker. Lane 1-7 represents annealing temperature 65.0℃, 63.8℃, 62.0℃, 59.1℃, 55.7℃, 52.9℃ and 51.0℃. (b) In-fusion cloning of *EjCDPK25* overexpression vector bacterial liquid PCR. Lane 1-5 represents five replicates.

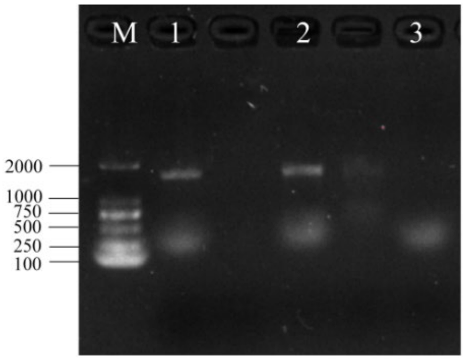


**Figure S2. Transgenic Arabidopsis T1 generation PCR identification target genes.**

*A. thaliana* transformed *EjCDPK25* over-expression vector PCR verification. M represents DL2000 DNA marker. Lan 1-3 represents three *A. thaliana* T1 generation line respectively.
